# PFRED: A computational platform for siRNA and antisense oligonucleotides design

**DOI:** 10.1101/2020.08.25.265983

**Authors:** Simone Sciabola, Hualin Xi, Dario Cruz, Qing Cao, Christine Lawrence, Tianhong Zhang, Sergio Rotstein, Jason D Hughes, Daniel R Caffrey, Robert Stanton

**Affiliations:** Medicinal Chemistry, Biogen, Cambridge, MA 02142, USA; Rgenta, Cambridge, MA, 02140, USA; Medicinal Chemistry, Ra Pharmaceuticals, Cambridge, MA, 02140, USA; Business Technology, Pfizer, Cambridge, MA 02140, USA; Computational Biology, Foundation Medicine, Cambridge, MA, 02141, USA; Department of Medicine, University of Massachusetts Medical School, Worcester, MA 01605, USA; Simulation and Modeling Sciences, Pfizer, Cambridge, MA 02139, USA; Chemical Engineering, Northeastern University, Boston, MA 02115, USA

## Abstract

PFRED a software application for the design, analysis, and visualization of antisense oligonucleotides and siRNA is described. The software provides an intuitive user-interface for scientists to design a library of siRNA or antisense oligonucleotides that target a specific gene of interest. Moreover, the tool facilitates the incorporation of various design criteria that have been shown to be important for stability and potency. PFRED has been made available as an open-source project so the code can be easily modified to address the future needs of the oligonucleotide research community. A compiled version is available for downloading at https://github.com/pfred/pfred-gui/releases as a java Jar file. The source code and the links for downloading the precompiled version can be found at https://github.com/pfred.

## Introduction

Over the last three decades small interfering ribonucleic acid (siRNA) and antisense oligonucelotides (ASO) have emerged as powerful tools to modulate the expression of genes in vivo and in vitro. Potential oligonucleotide therapeutics have been discovered and developed with mixed success[1,2]. For example, Mipomersen, Inotersen, Patisiran and Givlaari have all been approved for use in recent years. Mipomersen is a lipid lowering agent that targets the ApoB gene [3,4] which was eventually approved by the United States Food and Drug Administration (FDA) in 2013 and later discontinued due to adverse effects [5], Inotersen is an antisense compound targeting transthyretin (TTR) for the treatment of polyneuropathy caused by hereditary transthyretin-mediated amyloidosis [6], Patisirian was the first siRNA approved by FDA also targeting transthyretin-meidated amyloidosis [7], and Givlaari targets aminolevulinic acid synthase 1 (ALAS1) for the treatment of acute hepatic porphyria a rare genetic condition in which patients struggle to produce heme [8]. Furthermore, the approval of Nusinersen for the treatment of spinal muscular atrophy (SMA) in pediatric and adult patients represents a historical achievement for the SMA community and a success story for the field of nucleic acids based therapeutics [9].

However, the development of oligonucleotide therapeutics remains challenging as they often have unintended off-target effects and cause non-specific hepatic and renal toxicity. Additionally, it is exceptionally difficult to deliver oligonucleotides to most tissue types and organs. This can result in the need for tissue specific delivery systems and several siRNA compounds have advanced into development using this paradigm; these include liposomally liver targeted ALN-RSV01 which targets RSV through the nucleocapsid “N” gene [10], PF-4523655 for age related macular degeneration targeting the eye through direct injection [11] and galNac conjugates [12] such as the late stage ALN-AT3 [13] targeting antithrombin for the treatment of hemophilia. The recent FDA approval of Patisiran[14] represents a first-of-its-kind RNA interference (RNAi) therapeutic for the treatment of the polyneuropathy of hereditary transthyretin-mediated (hATTR) amyloidosis in adults and the first ever FDA approval of a siRNA treatment [7].

Designing antisense oligonucleotides (ASOs) and siRNA can be logistically challenging given the many competing design criteria that can be incorporated into the selection of tool or therapeutic sequences. Design considerations may include splice variants, cross-species targeting for validation, single nucleotide polymorphisms (SNPs), secondary structure, undesirable motifs (e.g. toxic, poly-A, or poly-G repeats), complementarity with off-target sequences, intron/exon boundaries, chemical modification pattern of nucleotides, and predicted activity. These factors need to be weighed according to the intended use of the oligonucleotide. For example, compounds designed as in vitro tools only need to be active in a single species, whereas it is advantageous for therapeutic oligonucleotides to be active in model organisms as well as patients. A number of computational tools have been developed to address different aspects of the design process, including OptiRNAi [15], siExplorer [16], DSIR [17], i-Score [17,18], MysiRNA-Designer [19], siDirect [20,21], siRNA Selection Server [22], siRNArules [23] and siRNA-Finder [24]. Additionally, major vendors of siRNA reagents such as Dharmacon (siDESIGN Center) [25], GeneScript (Target Finder) [26] and Thermo Fisher (BLOCK-iT) [27] have web based design tools available.

In the present paper we describe PFRED (**PF**izer **R**NAi **E**numeration and **D**esign tool), a client-server software system designed to assist with the entire oligonucleotide design process, starting with the specification of a target gene (Ensembl ID) and culminating in the design of siRNAs or RNase H-dependent antisense oligonucleotides. Sequences are chosen using bioinformatics algorithms built upon careful mining of the sequence-activity relationships found in public datasets as well as internal collections. The tool provides researchers with a user-friendly interface where the only required input is an accession number for the target gene and it returns a list of properties that are believed to contribute to the efficacy of an siRNA or ASO. These properties include human transcripts and cross-species homology, GC content, SNPs, intron-exon boundary, duplex thermodynamics, efficacy prediction score and off-target matches. An automated oligonucleotide selection procedure is available to quickly select one potential set of sequences with an appropriate property profile. The selection protocol can be customized by the user through changes of the selection cutoffs or the addition of alternate design parameters and algorithms.

## Materials and Methods

PFRED is implemented using a client-server software architecture. On the client-side, the graphical user interface (GUI) was developed in the Java programming language and is deployed to a user’s desktop through Java Web Start, the user will need to have a recent version of the Java development kit setup in their system. Because of the cross-platform support of Java, PFRED can be run from multiple desktop environments such as Windows, Mac and Linux. The most CPU-intensive algorithms used in PFRED are implemented as Python and Perl scripts which are wrapped within a docker container that works as a web service hosted in an Amazon Web Service (AWS) instance and invoked remotely from the client through a RESTful API, also developed in Java. Beyond the performance improvements, the client-server architecture also greatly simplifies the integration of disparate design algorithms that often rely on heterogeneous third-party libraries or reference datasets. Calls to the server-side algorithms are made through the PFRED java web service which uses the REST API that was developed using the Glassfish Jersey Java modules. New algorithms can be added to PFRED following the conventions established for the existing tool.

To facilitate the use of PFRED, users only need to download the latest GUI release and run it on their own computer; it is also possible for users to build their own private PFRED service by downloading the PFRED Docker repository in order to create and run the PFRED Service container following the documentation given at https://github.com/pfred/pfred-docker. This technique named containerization, facilitates cloud and server agnosticism, because the container automatically builds the needed environment for the PFRED back-end to work; users can also use this local service to test their own alternative algorithms or models, and deploy the PFRED service in a cloud environment like AWS, Azure, Google cloud, or on a local server. It is only required for the user to have Docker and Docker-compose setup in the system (see https://docs.docker.com/compose/). The current client-server architecture consists on an already built-in Docker Service within a publicly queryable AWS instance managed by Biogen, which offers a REST API for easy PFRED end-point access to all the back-end services (see https://github.com/pfred/pfred-rest-service for documentation on how to compile the REST source code and access the PFRED end-points using a browser). The PFRED GUI takes advantage of the RESTful API in order to access all the needed end-points for ASO and siRNA design (Schema 1 illustrates the PFRED architecture).

Figure 1 gives an example of the main PFRED interface showing the enumerated antisense sequences and data annotations related to these compounds. Oligonucleotide sequences and their properties can be easily visualized and analyzed in the main table. This main PFRED table provides most of the common spreadsheet functions such as alphanumeric sorting, column/row resizing, column coloring and conditional coloring. In addition, users can choose which properties are displayed in the table and customize the display order by drag-and-drop. To help users sift through large numbers of sequences, the main table supports filtering by either sequence patterns or property values. For example, a user can search for oligonucleotides that are conserved between human and mouse and have high predicted siRNA efficacy. Oligonucleotides can also be organized into different groups and then reviewed separately. Additional functions are also included to help users analyze their data. For example, the mathematical formula feature can be used to apply weights to multiple properties and summarize them into a single score for ranking and selection. The data can be exported as a comma separated file (CSV) and then imported into other applications for further analysis.

**Figure 1.**
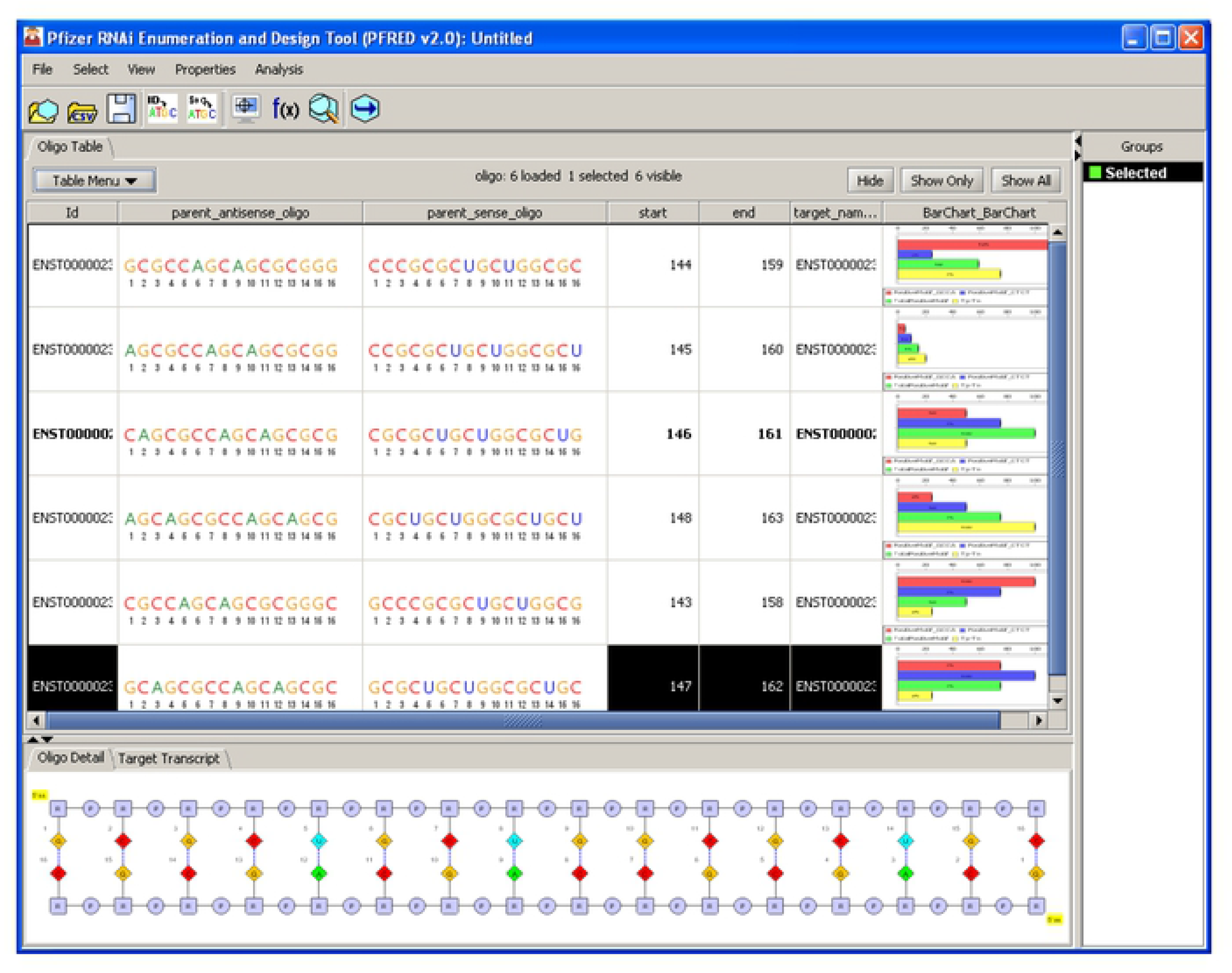
PFRED user interface. The main “Oligo Table” stores all the data generated through the oligo enumeration workflow (ol go sequences w th their properties) and provides access to common spreadsheet functions. It is possible to assign selected oligo sequences to specific groups which can be shown and accessed under “Groups” on the right-side panel. More detailed information about the oligo sequence notation and the primary mRNAtranscript can be visualized at the bottom of the user interface by enabling the “Oligo Detail” and “Target Transcript” panels.

A key element of PFRED’s functionality is its canonical representation of oligonucleotide structures. In a separate paper [28] we presented a monomer based notation language to represent complex biomolecule structures (the Hierarchical Editing Language for Macromolecules or HELM) and a desktop application named PME (Pfizer Macromolecule Editor) for their drawing and visualization, which has become an industry standard for biopolymer notations [29]. Figure 2A shows an example of HELM notation for a short antisense gapmer. As polymeric biomolecules, antisense oligonucleotides and siRNAs can be represented with HELM notation and different sequence visualization options can be created for specific purposes. Included within PFRED are three oligonucleotide representations, shown in Figure 2B-D. Figure 2B shows an explicit component or monomer view of the oligonucleotide breaking it down into sugars, bases and linkers. A sequence view is shown in Figure 2C, using a standard notation with RNA (capital) and DNA (lowercase) but emphasizing “chemical modifications” through the use of the pink dots. For example, the sequence shown in Figure 2C has a fully phosphorothioated backbone which is denoted by the pink dots between the base letters. It also has three locked nucleic acids (LNAs) at the 3’ and 5’ ends of the compound that are distinguished by the pink dots above the letters denoting these nucleosides. A final block representation is shown as Figure 2D. The block representation was created to emphasize patterns of chemical modifications to a structure. Base modifications are shown as pink dots below the blocks, sugars are colored depending on their structure (e.g. LNA-yellow, 2’OMe-Blue, etc.) and backbones other than phosphodiesters are represented by pink connectors above the blocks.

**Figure 2.**
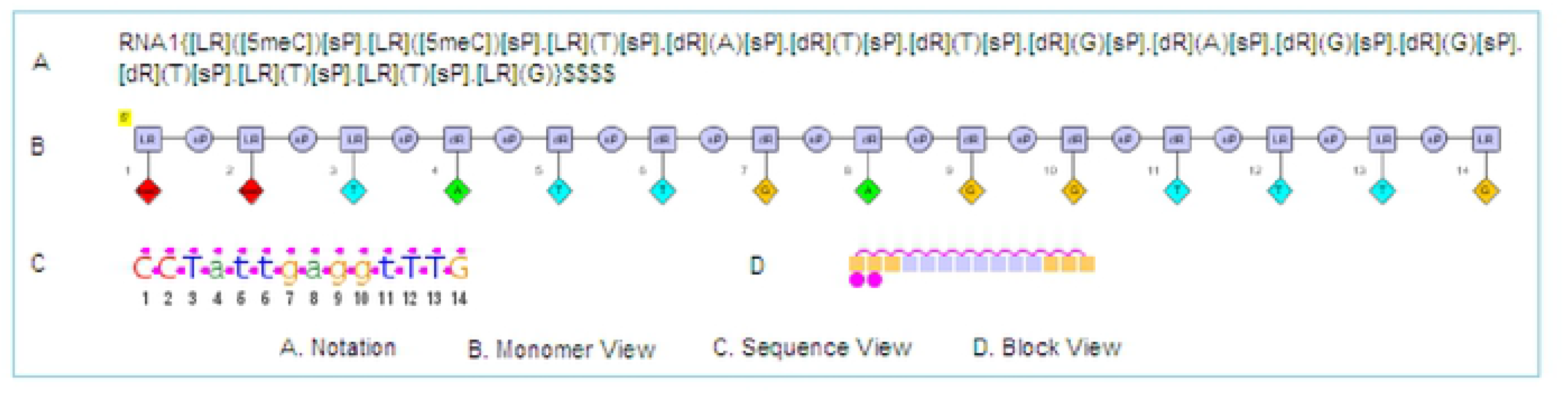
Alternative representations of ashort antisense gapmer sequence using the HELM notation (A) and different sequence visualizationoptions (B-D).

The PFRED enumeration workflow is shown in Figure 3. Transcript IDs and sequences of a target for different ortholog species (human, rat, mouse, dog, chimpanzee and macaque) are retrieved from the public ENSEMBL database through the ENSEMBL REST API (please refer to https://rest.ensembl.org/) given a single transcript or gene ID. Users define a primary transcript sequence of their length of choice to be enumerated (Figure 3, steps 1-3). Each enumerated oligonucleotide is then aligned to the other selected transcripts including splice variants and ortholog transcripts using the fast memory-efficient short read aligner, Bowtie, against the pre-calculated Bowtie index covering all transcript sequences [30]. Alignment results are processed by a Python interface and the number of mismatches to each transcript is reported back to PFRED as properties of the oligonucleotides. Exon location and SNP information are also retrieved through the EnsEMBL APIs for each base position and mapped to each oligonucleotide as properties.

**Figure 3.**
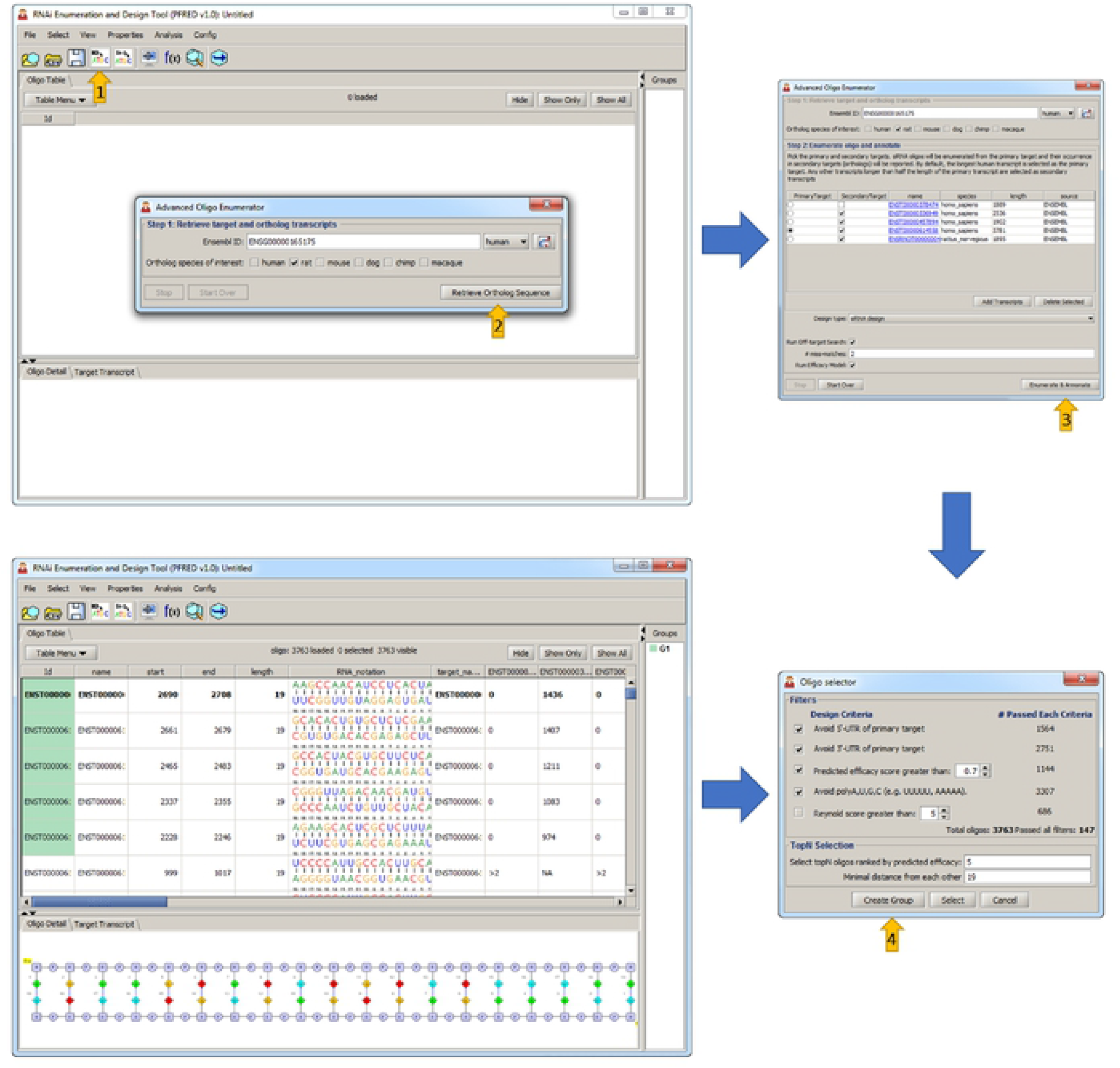
Oligo Enumeration and Annotation workflow. The process of designing an ASO or siRNA in PFRED starts with the Advanced Oligo Enumerator (**1**) by providing the Ensembl ID of a target of interest and the list of ortholog species to design against. The Retr eve Ortholog Sequence button (**2**) uses the Ensembl REST API to communicate with the public Ensembl database to retrieve all the sequence information related to the gene of interest (splice variants and ortholog transcrpts). At this point the user has the option to choose the Design type (siRNA design or AntiSense design) as well as additional ol go annotations (Offtarget and Efficacy predict ons) before tr ggering the Enumerate & Annotate button (**3**). Once all the descriptors have been calculated for each potential ASO or siRNA the design workflow becomes a process of selection (**4**).

Pre-built bowtie indexes for all species are required for off-target searches. These were built by creating a Python interface that automatically downloads all transcripts (cDNA) and unspliced gene sequences for any species in FASTA format using the ENSEMBL REST API which provides the advantage of always relying on the latest version of the ENSEMBL database. The FASTA files are written with transcript ID and Gene ID included in the sequence names. These FASTA files are used to build indexes for searching cDNA and gene off-target matches. The ENSEMBL IDs are used for post-processing the bowtie alignment results to calculate the number of genes that an oligonucleotide aligns to with 0, 1 and 2 mismatches, and to exclude the intended target from the off-target count. For siRNA, only the cDNA database is searched for both the guide and passenger strands. In contrast, for antisense design, only the sense sequences (or antisense sequence in reverse complementary direction) is searched against both the cDNA and unspliced gene sequence indexes for off-target hits. The number of cDNA off-target hits is then reported as an oligonucleotide property for 0, 1 and 2 mismatches.

Due to the cost of synthesizing and testing siRNAs and ASOs, rational design represents a critical step in any oligonucleotide experiment. A priori prediction of a compound’s function is a key component of RNAi and ASO applications to obtain effective silencing of the desired gene. Early models were based on empirical rules for designing functional siRNA, obtained from the experimental results of ASO and RNAi screening campaigns. For RNAi sequence design, Tuschl [31,32] first and later Reynolds [33] reported a series of guidelines for the implementation of a “point-based” scoring scheme where the presence or absence of key features in an siRNA duplex will lead to an increase or decrease of the final functional score. Several other algorithms have emerged since then [16,17,34–39] based on more sophisticated scoring schemes which use sequence descriptors and supervised machine learning techniques to select siRNAs capable of inducing effective gene silencing in cell lines. In PFRED, an algorithm for predicting siRNA functionality has been implemented by using diverse sets of oligonucleotide descriptors combined with a support vector machine (SVM) algorithm. Details of the sequence descriptors and algorithm implementation were previously described elsewhere [40]. Models of ASO activity have also improved but can be highly dependent on the length of the ASO as well as its chemical modification. Most published models use either sequence motif based filters [41] or computational approaches including secondary structure prediction [42–44], thermodynamics [45,46], machine learning [47–49] and molecular dynamics simulation [50], but all have reported only limited success. Here we integrated both the thermodynamics parameters calculated using Oligowalk [51] and the sequence motifs reported in Matveeva [41]. Using these parameters, a scoring scheme was derived that penalized motifs correlated with toxicity or promiscuity, strong inter- and intra-molecular binding energy as well as weak binders to the RNA target. The model was derived and tested using data from AOBase [52] including more than 500 ASOs against 46 targets. The thermodynamic calculations were based on unmodified DNA antisense oligonucleotides and therefore of limited relevance to modifications that significantly increase the melting temperature (Tm) of DNA/RNA duplex such as LNA.

Once all the descriptors have been calculated for each potential ASO or siRNA the design workflow becomes a process of selection. An example of the PFRED selection interface for siRNA design is shown in Figure 3, step 4. Oligonucleotide selections are made by first filtering out those compounds which align with: 1) known human SNPs, 2) sequence motifs associated with non-specific binding (such as polyG, polyT) and 3) low complexity sequences that have a high number of perfect off-target matches. An additional filter for ortholog transcript matches can be added depending on the experimental design requirement. Also, if predicted activities are available, they may be included as a selection criterion (models may depend on a specific oligonucleotide length or modification type). Other oligonucleotide properties such as number of off-target matches and diversity of location of each oligonucleotide along the gene sequence can be used to finalize the selection of desired compounds for a given experiment. The final selected compounds can then be exported in text format in preparation for synthesis or further analysis outside of the tool.

## Conclusions

We present here an informatics platform for the design of antisense and siRNA oligonucleotides. The system includes a basic set of design algorithms but is extensible such that experienced developers can add novel or proprietary descriptors. Workflows are included which walk researchers through the process of identifying a target sequence and choosing the design criteria to be applied in selection of candidate molecules. A series of oligonucleotide visualization and plotting algorithms are included along with spreadsheet functions allowing data import/export, formulas, and conditional formatting. Links to the source code and hosted versions can be found at https://github.com/pfred.

## Aknowledgements

The authors would like to thank Art Kreig, Matteo Di Tommaso and Enoch Huang for their support of this work. Dario Cruz acknowledges support from the Biogen Internship & Co-op program.

---

**Schema 1.**
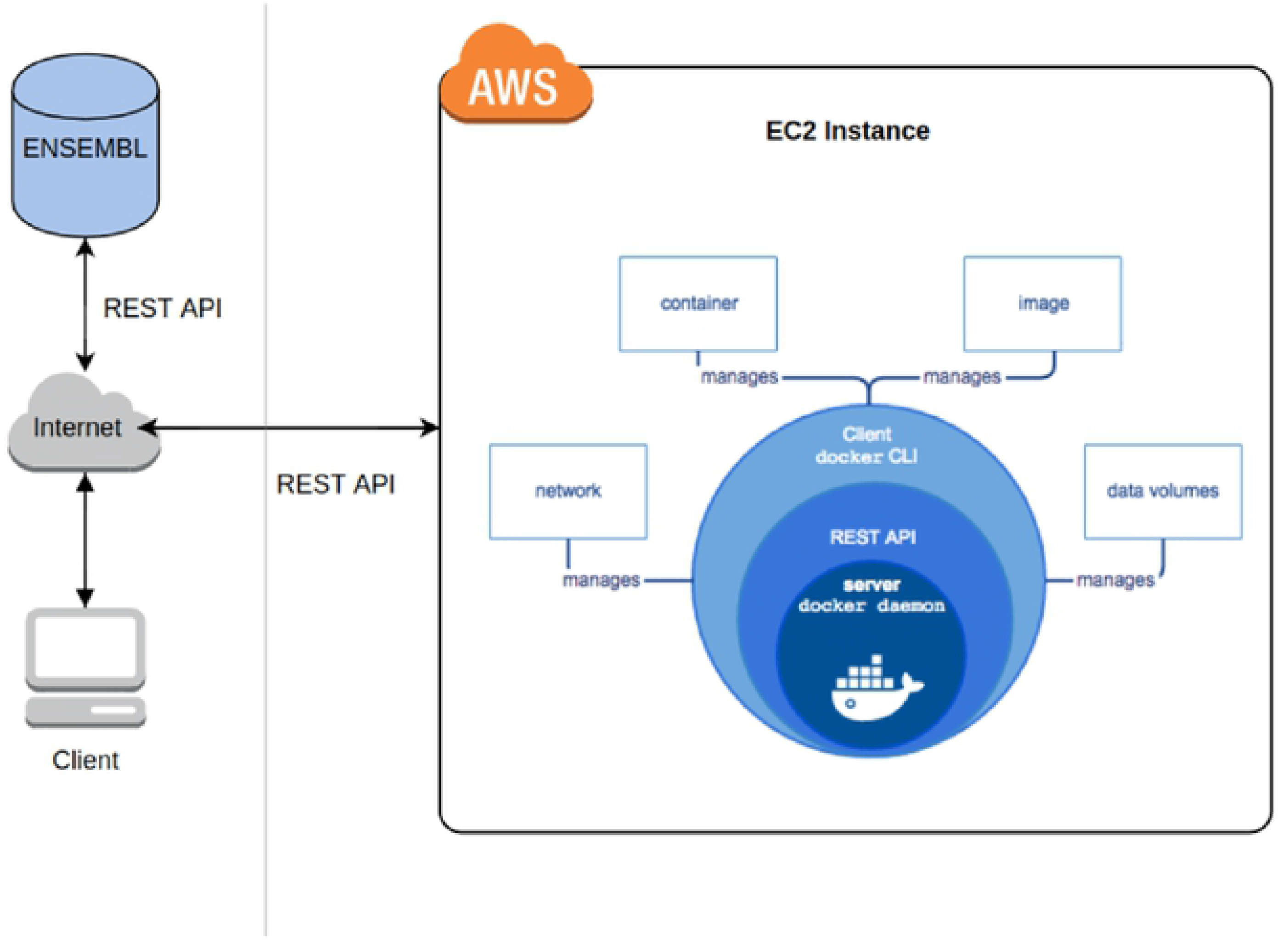
PFRED client server architecture based on the AWS and Docker containerization.

